# *LincRNA-Cox2* regulates smoke-induced inflammation in murine macrophages

**DOI:** 10.1101/2022.07.28.501765

**Authors:** Mays Mohammed Salih, Elektra Kantzari Robinson, Eric Malekos, Elizabeth Perez, Allyson Capili, Kihwan Kim, William Z. Zhang, Suzanne M. Cloonan, Susan Carpenter

## Abstract

Cigarette smoke (CS) exposure is a risk factor for many chronic diseases including chronic obstructive pulmonary disease (COPD), however the mechanism by which smoke exposure can alter homeostasis and bring about chronic inflammation is poorly understood. Here, we showcase a novel role for smoke in regulating long noncoding RNAs (lncRNAs), showing that it activates *lincRNA-Cox2*, which we previously characterized as functional in inflammatory regulation. Exposing *lincRNA-Cox2* murine models to smoke *in vivo* confirmed *lincRNA-Cox2* as a regulator of inflammatory gene expression in response to smoke both systemically and within the lung. We also report that *lincRNA-Cox2* negatively regulates genes in smoked bone marrow derived macrophages exposed to LPS stimulation. In addition to the effects on lncRNAs, we also report dysregulated transcription and splicing of inflammatory protein-coding genes in the bone marrow niche following CS exposure *in vivo*. Collectively, this work provides insights into how innate immune signaling from gene expression to splicing is altered following *in vivo* exposure to CS and highlights an important new role for *lincRNA-Cox2* in regulating immune genes following smoke exposure.

## Introduction

Chronic obstructive pulmonary disease (COPD) is a debilitating inflammatory lung disease associated with tobacco smoking. The pathogenesis of COPD remains poorly understood but is known to involve abnormal inflammatory responses of lung macrophages that are induced by cigarette smoke (CS) exposure (1). Macrophages specialize in the initial response to external signals such as infectious agents, environmental pollutants as well as tissue damage (2). CS exposure can attenuate macrophage function including altering responses to infectious agents (2) and suppressing phagocytic rate, (3) in addition to increasing pro-inflammatory cytokine and chemokine production (4,5). CS-driven lung inflammation is also linked to tissue injury, susceptibility to microbial infections, and poor wound healing (6,7). The mechanism by which CS contributes to the heightened inflammation observed in COPD is not well understood.

Alternative splicing is a key regulator of inflammation and a potential mechanism that links CS exposure to abnormal inflammatory response in macrophages in COPD (8). Alternative splicing is a highly regulated process which allows for multiple isoforms of a protein to be produced from a single gene (9-12). Previous work identified important splicing events in genes involved in chemokine signaling and metabolism following lung inflammation (13,14). With relevance to COPD, comparisons of bronchial biopsies of smokers versus non-smokers suggest that smoke induces alternative splicing of the receptor for advanced glycation end products (AGER or RAGE), which in turn is known to have a role in mediating lung endothelial inflammation in COPD (15). However, a robust understanding of how CS can alter gene expression and isoform usage specifically in bone marrow-derived macrophages (BMDMs) has not been investigated. Here we made use of multiple splicing tools and identified retained intron events as the top alternative splicing events occurring in macrophages following *in vivo* smoke exposure.

To better understand CS induced transcriptional modulation, we also focused on expression changes in long noncoding RNA (LncRNA) genes in addition to coding genes. LncRNAs have emerged as key regulators of gene expression and as effective modulators of immune cell development and function (16,17). We have previously characterized the function of the inflammatory inducible *lincRNA-Cox2* using both *in vitro* and *in vivo* approaches where it plays a broad regulatory role during inflammation (18, 19). *LincRNA-Cox2* can function in *cis* through an RNA enhancer mechanism to regulate its neighboring protein Ptgs2 (Cox2), as well as functioning in *trans* to globally regulate expression of innate immune genes (19-26). In this study, we identified *lincRNA-Cox2* as one of the most upregulated genes in macrophages following *in vivo* smoke exposure. We used two different *lincRNA-Cox2* mouse models to determine the effect of modulating *lincRNA-Cox2* expression on CS-induced macrophage inflammatory response. Our *lincRNA-Cox2* deficient (mutant, Mut) model represents *lincRNA-Cox2* knockdown where the splice sites have been targeted with CRISPR/Cas9 as previously reported (18). In addition, we used our recently generated rescue transgenic mouse model that involved crossing the *lincRNA-Cox2* transgenic overexpressing line with our *lincRNA-Cox2* deficient (Mut) line labeled MutxTg (19). The two models allow to us to both investigate the *trans-*regulatory role for *lincRNA-Cox2* independent of its *cis* effects on its neighboring gene Ptgs2 and whether ubiquitous overexpression of *lincRNA-Cox2* from a non-native locus could rescue the phenotypes we observed in the *lincRNA-Cox2* deficient mice. Utilizing a physiologically relevant model of *in vivo* smoke exposure, we found that *lincRNA-Cox2* functions in the lung to regulate inflammatory gene expression following CS exposure, and that it can regulate inflammatory genes in *trans* under these conditions.

## Results

### *In vivo* CS exposure alters the expression of genes at the RNA and protein levels in BMDMs

To determine the impact of CS exposure on inflammatory pathways, we utilized a well-established CS-induced COPD model, whereby WT mice were subjected to main and side stream CS for 8 weeks (27-30). Bone marrow from room air (RA) and CS exposed mice was harvested and differentiated into bone marrow-derived macrophages by culturing for 8 days in L929 media. RNA sequencing (RNA-seq) was performed to investigate the effect of *in vivo* CS exposure on the macrophage transcriptome (**Fig. 1A**). Differential expression (DESeq2) analysis revealed a significant number of dysregulated protein-coding genes following CS exposure (**Fig. 1B**). Interestingly, most protein coding genes were up-regulated (102 genes) while only 22 protein coding genes were down-regulated following CS exposure. To investigate if there was a common regulatory mechanism controlling the up-regulated genes, we used “i-cis target” (31), an analysis software that determines enriched transcription factor (TF) motifs (**Fig. 1C**). I-cis target showed high enrichment scores for inflammatory response related TFs, including type I IFN response (STAT1, 3, and 4) and NF-kB pathway (Rela) (**Fig. 1C**). Of all the TFs associated with CS exposure, only Cebpb and Cepbd were found to be significantly upregulated at the transcriptional level (**SFig. 1**). To identify biological processes and pathways impacted by CS, we performed gene ontology (GO-term) analysis on up and down-regulated protein coding genes which, revealed enrichment for immune response associated pathways, including Il1, Ifng, and Tnf response biological processes as well as the involvement of NF-kB and JAK-STAT in KEGG pathways (**Fig. 1D, E**). We also observed enrichment of KEGG pathway terms associated with hypoxia as well as inflammatory diseases such as rheumatoid arthritis and cancer, both of which are associated with CS-exposure (**Fig. 1D**) (32, 33). GO-term analysis of downregulated protein-coding genes was enriched in the regulation of rho protein signal transduction (**SFig. 2**), previously reported to be dysregulated by smoke exposure (34, 35). To determine whether CS exposure effects extend beyond changes at the level of transcription, we next measured cytokine and chemokine protein production changes secreted from CS-exposed BMDMs by ELISA, finding that Il9, Il16, Gcsf, and Vegf were all significantly downregulated by CS (**Fig.1F-I**). These results revealed that CS can shape the bone marrow niche, impacting a newly developed macrophage’s baseline levels of RNA and protein expression.

**Figure 1:**
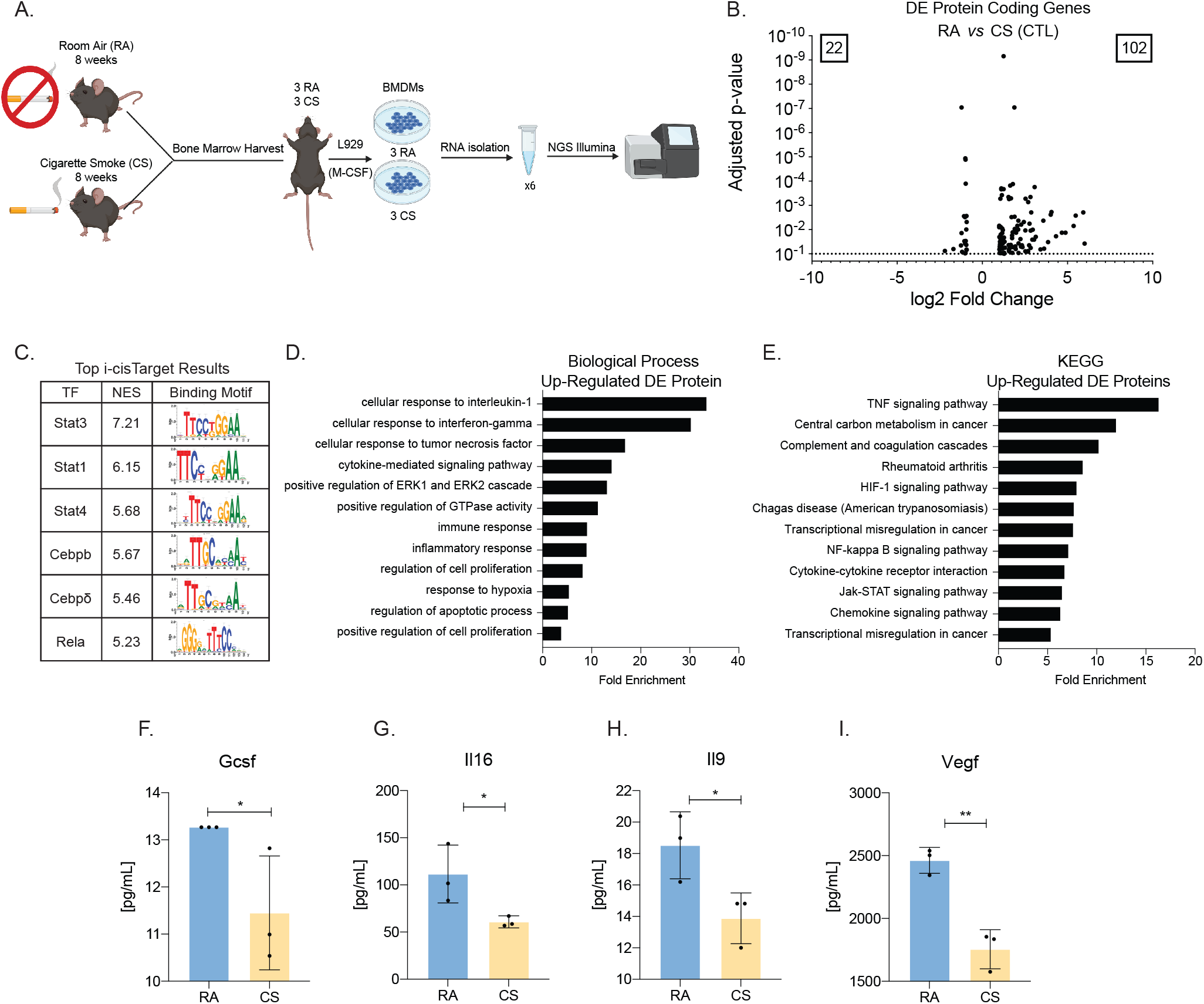
Cigarette smoke exposure activates inflammatory protein-coding genes in bone marrow-derived macrophages. (A) Schematic of cigarette smoke exposure and RNA-sequencing experiment L929 (MCSF) induced BMDMs. (B) Volcano plot of all differentially expressed protein-coding genes when comparing BMDMs from room air (RA) to cigarette smoke (CS) mice. (C) All promoters of upregulated protein-coding genes were analyzed using i-cisTarget to assess the top transcription factors regulating genes, sorted by normalized enrichment score (NES) and binding motif. (D) The associated biological process of upregulating differentially expressed protein-coding genes from CS exposure using DAVID tools. (E) The associated KEGG pathways of upregulated differentially expressed protein-coding genes from CS exposure using DAVID tools. The supernatant was harvested from RA and CS cultured BMDMs and multiplex cytokine analysis was performed for (F) Gcsf, (G) Il16, (H) Il9, (I) Vegf. Each dot represents an individual animal. Error bars represent the standard deviation of biological triplicates. Student’s t-tests were performed using GraphPad Prism. Asterisks indicate statistically significant differences between mouse lines (*p ≥ 0.05, **p ≥ 0.01).

### *In vivo* CS exposure drives alternative splicing events including intron retention in BMDMs

RNA-seq data is commonly used to determine changes in gene expression under different conditions, but it can also be used to profile alternative splicing events and uncover changes in isoform usage. To this end we utilized two different splicing tool pipelines, rMATS and JuncBASE+DRIMSeq (**SFig. 3**). We incorporated information from both toolsets and performed manual inspection of reported splice events via sashimi plots. Here we showed that most genes undergoing alternative splicing following CS exposure were not differentially expressed (**Fig. 2A, SFig. 4**). In total, rMATS identified 425 splicing events while JuncBASE+DRIMSeq identified 47 that occur in macrophages following *in vivo* CS exposure (padj < 0.1 |ΔPSI| > 10) (**Fig. 2B**). The dominant alternative splicing events we identified using both JuncBASE and rMATS were retained intron (RI) events (**Fig. 2B**), with a total of six RI events across four genes common to both JuncBASE and rMATS analysis (**Fig. 2C**). One prominent RI event was preferential retention in the RA condition of the intron between exons 11 and 12 (as annotated in Gencode vM18 basic) of transcripts coding Mib2, a ubiquitin-protein ligase that mediates ubiquitination of proteins in the Notch signaling pathway (**Fig. 2D, SFig. 5**). We validated this event using RT-PCR, confirming a significant decrease in the retained intron isoform under CS conditions (**Fig. 2D-F, SFig. 6**). These results highlight how CS exposure can modify the macrophage transcriptome by ways other than gene expression changes and showcases alternative splicing as a mechanism for functional modulation post smoke exposure.

**Figure 2:**
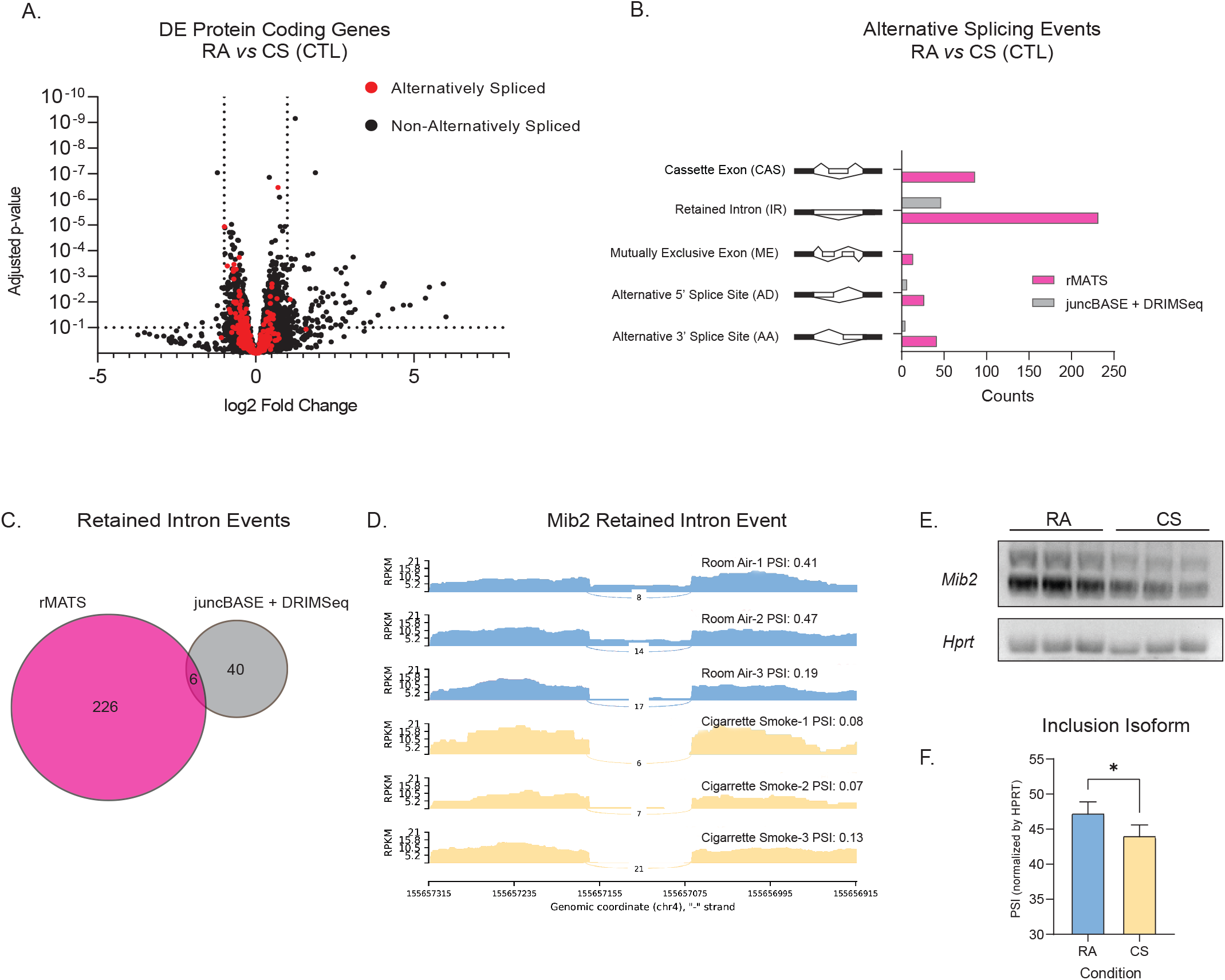
Cigarette Smoke COPD induces alternative splicing in bone marrow-derived macrophages. (A) Significant event counts from rMATS and JuncBASE+DRIMSeq. Events were considered significant if FDR < 0.1, |PSI| > 10. (B) Venn diagram of significant retained intron events w^1^ith shared events determined by Bedtools’ intersect function. (C) Sashimi plot of the Mib2 retained intron event. (D) RT-PCR gel results of Mib2 at the event site and loading control HPRT in biological triplicates and (E) percent spliced in (PSI) as determined by the average relative intensity of the inclusion form bands. Error bars represent standard deviation of biological triplicates. Asterisk indicates statistically significant differences between conditions using Welch’s t-tests (*p < 0.05).

### LncRNAs are differentially expressed in response to CS exposure in BMDMs

Since CS exposure impacts transcription and splicing of protein-coding genes, we next determined if CS also influenced the expression of lncRNAs, which are themselves often products of alternative splicing. Using DESeq2 we identified 47 lncRNAs that were downregulated and 3 that were upregulated following *in vivo* CS exposure **(Fig. 3A**). LncRNAs can function in *cis* to regulate their neighboring genes or in *trans* to regulate genes on different chromosomes (16). To gain insight into the mechanism of action of the dysregulated lncRNAs, we first performed bioinformatic analysis to determine the expression of the proteins neighboring the lncRNAs to ascertain if they are co-regulated which can provide insights into the mechanism of action of the lncRNAs (guilt by association) (36). We utilized the genomic regions enrichment of annotation tool (GREAT) (37) to identify the neighboring protein-coding genes for all down-regulated lncRNAs based on the genomic location (**Fig.3B**). Gene ontology (GO) analysis of the proximal protein-coding genes are associated include negative regulation of Tgfb production and mRNA splicing regulation (**Fig. 3C**). To further assess the guilt by association model and the possibility that these lncRNAs might function in a *cis-*regulatory mechanism, we examined the normalized counts of the neighboring protein-coding genes in our RNA-seq data and found that 9 of the protein-coding genes neighboring down-regulated lncRNAs were significantly differentially expressed. Specifically, 7 of the neighboring protein-coding genes were co-regulated while 2 were anti-correlated in expression (**Fig. 3D**). Of the 3 up-regulated lncRNAs all neighboring protein-coding genes were co-regulated (**Fig. 3E**). Most importantly, *lincRNA-Cox2* was identified as a significantly induced lncRNA following smoke exposure, similar to its neighboring protein-coding gene *Ptgs2* (*Cox2*), a critical inflammatory response gene (**Fig.3E, F**). These results highlight how CS exposure can impact the expression of lncRNAs that play important roles within the regulatory network that governs the innate immune response in macrophages.

**Figure 3:**
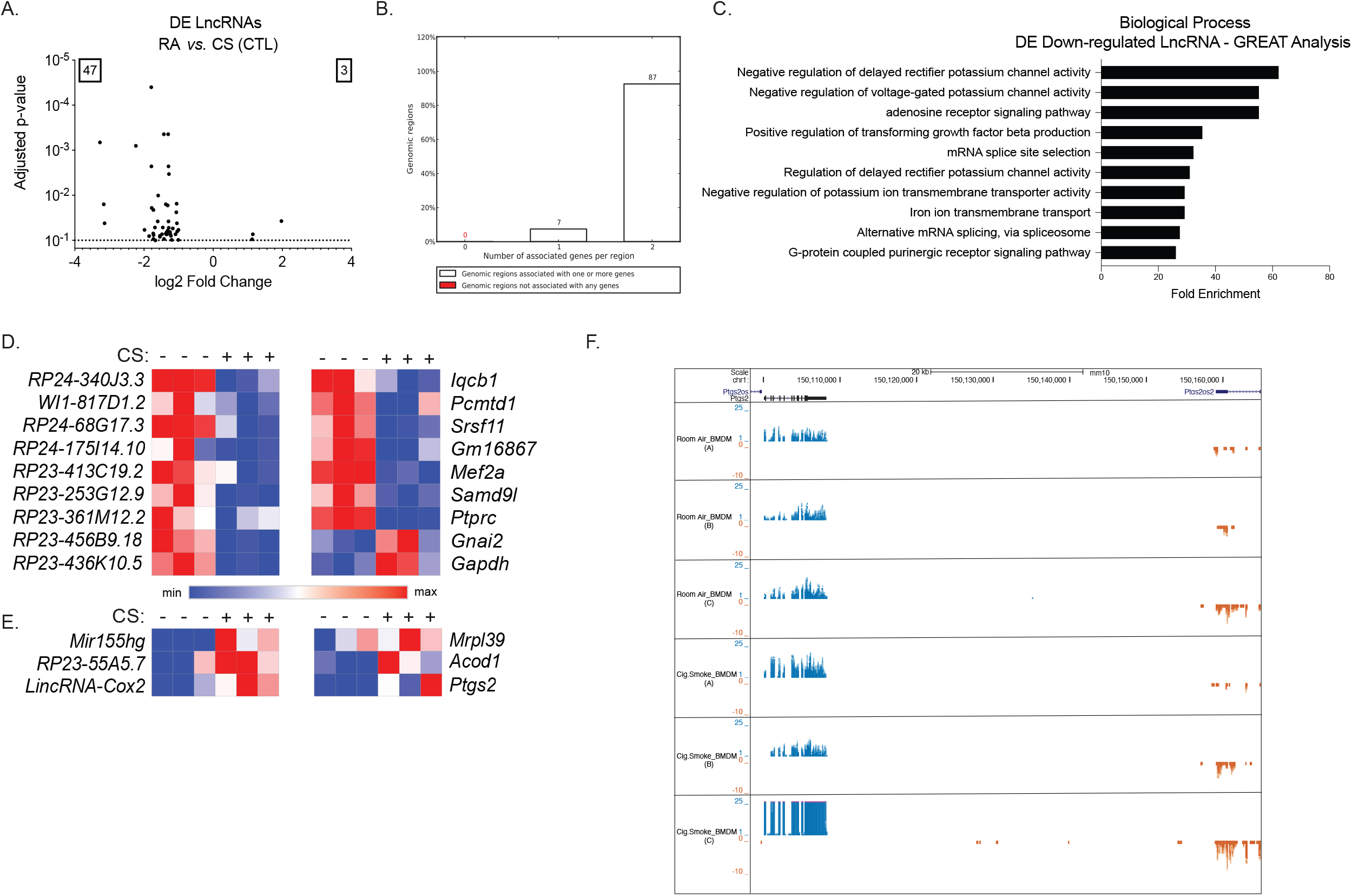
LncRNA regulated by cigarette smoke in bone-marrow-derived macrophages harvested from COPD mice. (A) Volcano plot of all differentially expressed long non-coding RNA genes when comparing BMDMs from room air (RA) to cigarette smoke (CS) mice. (B) Bar-graph of the number of associated genes per region generated using GREAT. (C) The neighboring protein-coding genes of upregulated differentially expressed lncRNA genes from CS exposure also determined the associated biological process using GREAT analysis. (D) A heat map of only significantly DE protein-coding and associated lncRNA genes from GREAT was generated with normalized read counts from RNA-seq of BMDMs with and without cigarette smoke (CS). (E) A heat map of up-regulated lncRNAs and associated protein-coding genes based on normalized read counts. (F) UCSC Genome Browser shot of 7 separate tracks. The top track is of M18 gene annotations, including Ptgs2 (black) and lincRNA-Cox2 (blue). Tracks following 2-4 tracks are stranded RNA-Sequencing reads of BMDM from room air (RA) mice. Tracks 5-7 are of stranded RNA-Sequencing reads of BMDM from cigarette smoked (CS) mice. Blue reads are positive stranded and the orange reads are negative stranded.

### Loss of *lincRNA-Cox2* results in dysregulated responses to CS exposure *in vivo*

We have recently reported that *lincRNA-Cox2* is a critical regulator of lung homeostasis and acute lung inflammation (19) but have yet to determine the impact of *lincRNA-Cox2* during smoke-induced chronic inflammation. Since *lincRNA-Cox2* is upregulated following CS exposure, we examined WT and *lincRNA-Cox2* deficient (Mut) mice (18) in an 8-week CS exposure model to explore the role for *lincRNA-Cox2*-mediated regulation in COPD pathogenesis (**Fig. 4A**). To assess CS-associated inflammation, we harvested bronchoalveolar lavage fluid (BALF), lung tissue, and serum of mice exposed to 8 weeks of RA or CS and measured cytokine and chemokine levels. These experiments revealed a disruption of cytokine and chemokine expression locally in the lung (from BALF and tissue) and globally from the peripheral serum (**Fig. 4 B-O**) in *lincRNA-Cox2* deficient mice. At baseline in the room air mice, we see that loss of *lincRNA-Cox2* results in an increase in Mip2, Il1a and Lix in the lung (**Fig. 4 N-O**) and a decreased in Il16 in the serum (**Fig. 4 G**). Both Gcsf and Il16 are known mediators in COPD pathogenesis (38, 39), and they were significantly downregulated in the BALF (**Fig. 4B-C**) and serum (**Fig. 4F-G**) while increased in the lung (**Fig 4J-K**) of *lincRNA-Cox2* deficient (Mut) mice. Moreover, we found that some dysregulated cytokines were localized to the BALF (**Fig. 4D-E**), serum (**Fig. 4H-I, SFig. 7B-C**), or lung tissue compartments (**Fig. 4L-O, SFig. 7D-I**). For instance, we see Mip2 (Cxcl2) is downregulated in the *lincRNA-Cox2* deficient mice only in BALF after CS exposure (**Fig. 4D**). Fractalkine is downregulated in BALF (**Fig. 4E**) and lung (**Fig.4M**), while Il1a is upregulated in the serum (**Fig. 4I**) and lung (**Fig. 4N**) of *lincRNA-Cox2* deficient mice. These data suggest that *lincRNA-Cox2* exerts different effects within the lung compared to the periphery, and that *lincRNA-Cox2* exert control beyond the transcriptional level and can modulate smoke-induced signaling at the protein level within the lung.

**Figure 4:**
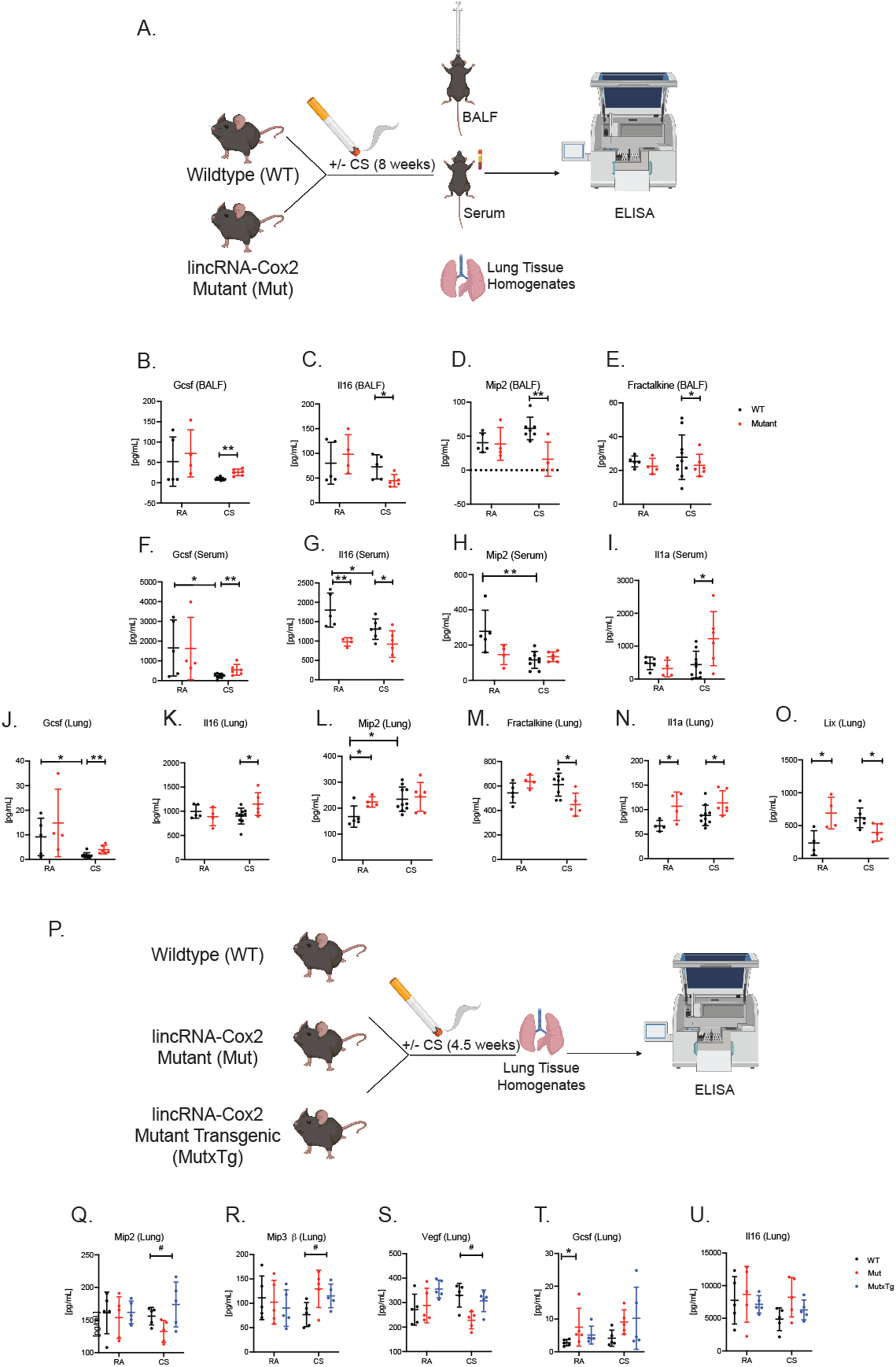
lincRNA-Cox2 positively and negatively regulates cytokines in a cigarette-smoke COPD model. (A) Experimental schematic of 8-week cigarette smoke COPD model using WT and lincRNA-Cox2 mutant mice. After 8 weeks BALF, serm and lung tissue were harvested for subsequent cytokine analysis. ELISAs were performed on room air and cigarette-smoked BALF samples for (B) Gcsf, (C) Il16, (D) Mip2 and (E) Fractalkine. Cytokine analysis was performed on serum for (F) Gcsf, (G) Il16, (F) Mip2 and (I) Il1a. ELISAs were also performed on normalized lung tissue samples for (J) Gcsf, (K) Il16, (L) Mip2, (M) Fractalkine, (N) Il1a and (O) Lix respectively from lung samples. (P) Experimental schematic of cigarette smoke COPD 4.5 week model for WT, lincRNA-Cox2 mutant and lincRNA-Cox2 MutxTg mice. Lung tissue was harvested for cytokine analysis to measure (Q) Mip2, (R) Mip3β, (S) Vegf, (T) Gcsf and (U) Il16 respectively. Each dot represents an individual animal. Error bars represent standard deviation of biological replicates. Asterisks indicate statistically significant differences between mouse lines using Student’s t-tests (*p < 0.05, **p < 0.01, ***p < 0.001). Student’s t tests were performed using GraphPad Prism to obtain p values.

### *LincRNA-Cox2* functions *in trans* to regulate gene expression

To gain insights into how *lincRNA-Cox2* might function mechanistically to control these gene expression changes, we wanted to determine if it was mediating its effects in *trans* since many of the affected cytokines are encoded on different chromosomes. To answer this question, we utilized our newly generated Mutant/Transgenic mouse model (MutxTg), whereby a transgenic mouse overexpressing *lincRNA-Cox2* ubiquitously in locus H11 was crossed with our *lincRNA-Cox2* deficient (Mut) mouse model (19). The hypothesis was that if *lincRNA-Cox2* is functioning in *trans* to regulate gene expression then the differences observed in the *lincRNA-Cox2* deficient (Mut) mice would be rescued by the overexpression of *lincRNA-Cox2* in the Mutant/Transgenic mouse model (MutxTg). We performed a 4.5-week CS-exposure COPD model on WT, *lincRNA-Cox2* deficient and *lincRNA-Cox2* MutxTg, lung tissue was harvested to measure cytokine and chemokine expression (**Fig. 4P**). Interestingly, we found several cytokines and chemokines that are dysregulated in the *lincRNA-Cox2* mutant mouse and rescued in the *lincRNA-Cox2* MutxTg, including Mip2, Mip3b and Vegf (**Fig. 4Q-S**). Moreover, we found that while Gcsf and Il16 are not significantly differentially expressed at the 4.5-week timepoint between the *lincRNA-Cox2* deficient and the *lincRNA-Cox2* MutxTg mice, they trend upwards in the *lincRNA-Cox2* deficient mice and are unchanged comparing WT and MutxTg mice (**Fig. 4S-T**). In total these data suggest that *lincRNA-Cox2* functions *in trans* to regulate immune genes within the lung following CS exposure.

### *LincRNA-Cox2* regulates the acute inflammatory response in BMDMs pre-exposed to CS

Given the dysregulation of inflammatory inducible genes observed in BMDMs from CS exposure at resting states (**Fig. 1**), we next aimed to determine how macrophage inflammatory response to stimulus could be altered following CS exposure. To assess this globally, we isolated and generated BMDMs from CS exposed mice (8 weeks) and compared control to LPS stimulated cells (**Fig. 5A**). Overall, we observed a subtle ablated inflammatory response in protein-coding genes as well as an overall decrease in lncRNA genes response following *in vivo* CS exposure (**Fig. 5B-C**). For the differentially expressed lncRNAs we did not observe any co-regulation between them and their neighboring protein coding genes (**Fig. 5C**). We also performed ELISAs to measure protein production levels in LPS stimulated BMDMs that were exposed to CS *in vivo*, we found that Il16 and Il12 (p40) were downregulated following CS exposure while Mip3b (Ccl19) and Mcp-5 (Ccl12) were upregulated (**Fig. 5D-G**). These data emphasize the complexity of the signaling pathways that are impacted following *in vivo* continual exposure to CS.

**Figure 5:**
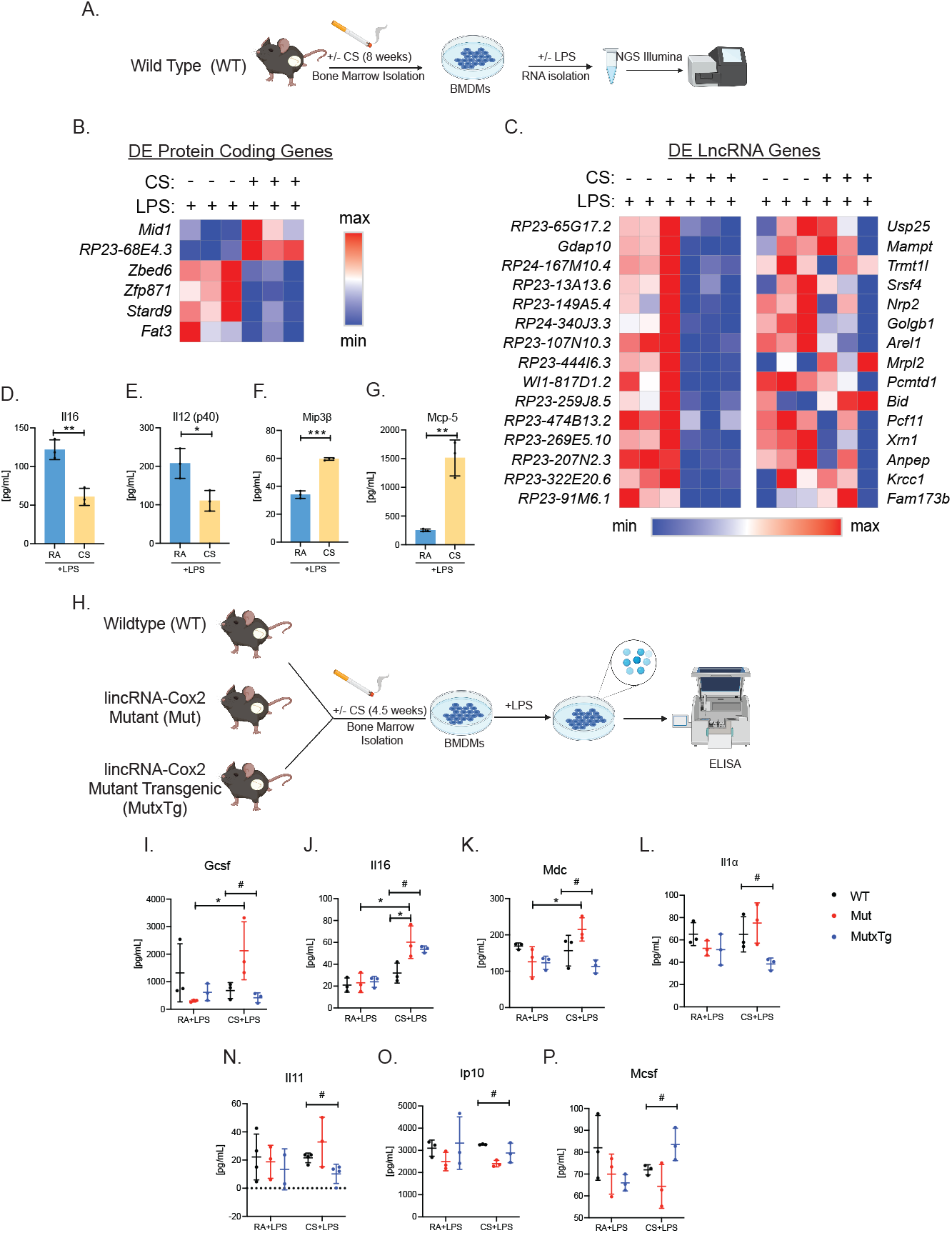
lincRNA-Cox2 regulates the impact of cigarette smoke during acute inflammation in bone marrow-derived macrophages. (A) RNA-sequencing experimental schematic of BMDM COPD with LPS. (B) Heatmap generated for normalized counts of protein-coding genes from room air and cigarette smoked BMDMs post LPS treatment. (C) Heatmap of lncRNA genes and their respective neighboring protein-coding genes normalized counts from room air and cigarette smoked BMDMs post LPS treatment. Cytokines measured from supernatants of BMDMs using ELISA technique for (D) Il16, (E) Il12(p40), (F) Mip3β and (G) Mcp5. (H) Experimental schematic for LPS treated BMDM harvest from WT, lincRNA-Cox2 mutant and lincRNA-Cox2 MutxTg mice exposed to room air or cigarette-smoked. Supernatants were harvested and ELISAs were performed to measure (I) Gcsf, (J) Il16, (K) Mdc, (L) Il1a, (N) Il11, (O) Ip10 and (P) Mcsf. Each dot represents an individual animal. Error bars represent standard deviation of biological triplicates. Asterisks indicate statistically significant differences between mouse lines using Student’s t-tests (*p < 0.05, **p < 0.01). Number sign indicate statistically significant differences between mouse lines using one-way Anova statistical test (#p<0.05). Student’s t tests and ANOVA tests were performed using GraphPad Prism to obtain p values.

*LincRNA-Cox2* has been widely studied in macrophages and shown to act as both a positive and negative regulator of immune genes following inflammatory activation (19-26). While it is clear *lincRNA-Cox2* provides a layer of regulation for inflammation driven by CS exposure *in vivo* (**Fig. 4**), we next aimed to elucidate how *lincRNA-Cox2* impacts the acute inflammatory response in BMDMs, from mice exposed to CS for 4.5 weeks by measuring cytokine and chemokine production using ELISAs (**Fig. 5H**). First, we found that BMDMs differentiated from 4.5-week CS exposed WT, *lincRNA-Cox2* deficient and *lincRNA-Cox2* MutxTg mice displayed no significant changes in cytokine levels between the genotypes either after RA or CS (**SFig. 8B-J**). However, we found lipopolysaccharide (LPS) treated BMDMs from *lincRNA-Cox2* deficient mice exposed to CS for 4.5 weeks exhibited higher Il16, Gcsf, Il1a, Mdc and Il11 in comparison to WT mice (**Fig.5 I-N**). Additionally, we found that Ip10 (Cxcl10) and Mcsf (Csf1) were downregulated in BMDMs from *lincRNA-Cox2* deficient mice (**Fig. 5 O-P**) compared to WT mice. Importantly, we found that all of the affected cytokines were rescued in the MutxTg mice, indicating that *lincRNA-Cox2* regulates these genes in *trans* through a yet unknown mechanism (**Fig. 5 O-P**). Overall, our data supports *lincRNA-Cox2’s* involvement in regulating the macrophage response to smoke exposure and its contribution to driving an exacerbated inflammatory response to stimulus in macrophages.

## Discussion

Cigarette smoking is a major risk factor for COPD which is characterized by chronic inflammation in the airways that results in irreversible airflow obstruction (40). In this study, we investigated the impact of smoke on the bone marrow (BM) niche using an *in vivo* murine smoking model (8). We were intrigued to find that BMDMs from 8-week CS exposed mice had a significant impact on the transcriptome. Transcriptional changes were not limited to gene expression; we showed that smoke induced signaling can modulate the splicing landscape of BMDMs and identified a prevalence of retained intron events in macrophages from smoked mice in addition to other events such as exon skipping (cassette exon). Interestingly, many of the proteins shown here to undergo alternative splicing are not themselves altered at the gene expression level in response to smoke exposure, highlighting the importance of splicing analysis to uncover changes to the transcriptome that gene expression changes studies cannot detect.

Our differential gene expression analysis revealed disruption in many proteins with many of them being overexpressed following smoke exposure and regulated by transcription factors such as STAT1, STAT3 and STAT4, as well as Cebpb, Cebpd, and NF-kB (41). Mechanistically, smoke induces oxidative stress in the lung which is thought to activate redox sensitive transcription factors (TFs), such as NF-kB, leading to inflammatory pathway activation (41). Additionally, STAT3 and STAT4 are also reported to be activated by smoke exposure and STAT3 was shown to impact immune cell recruitment and inflammatory cytokine production following smoke exposure (42, 43). While it is not surprising that many of the protein coding genes activated by CS belong to pathways that are under the regulation of NF-kB and STATs, our study is one of first to demonstrate this is occurring in the bone marrow compartment, a site distant from immediate smoke exposure. We confirmed this immune dysregulation at the protein level showing altered levels of Il9, Il16, Gcsf, and Vegf, all of which have been reported to be disrupted in response to CS exposure (38, 44-49). Trends of dysregulation (up or downregulation) of these cytokines following CS exposure are complex and appear in some cases to be cell type specific as well as being dependent on the nature and duration of exposure.

A recent study has reported that lncRNAs can be activated or repressed post long-term smoke exposure (50). Interestingly, we found an overall repression of macrophage lncRNAs and their associated protein coding genes in response to 8-week CS exposure, with *LincRNA-Cox2* being only one of three lncRNAs that were transcriptionally induced by CS. LncRNAs can serve as regulatory elements, modulating expression of either neighboring protein coding genes (*in cis*) or genes on distal chromosomes (*in trans*) (16,51). Specifically, we show that induction of *lincRNA-Cox2* is accompanied by induction of associated protein coding gene *Ptgs2* (*Cox2*), which is consistent with our previous findings (18). Most importantly, several lncRNAs that were upregulated, or downregulated following CS exposure showed similar patterns of expression compared to their neighboring protein coding genes, suggesting that they are co-regulated. For this study we focused our efforts to understand how *lincRNA-Cox2* is functioning following CS exposure, but future work could focus on the other 11 lncRNAs, their neighboring protein coding genes and their regulatory roles following CS exposure.

Since *lincRNA-Cox2* is induced by CS exposure, we wanted to determine if it plays any mechanistic role in regulating immune gene expression following CS. To this end, we exposed *lincRNA-Cox2* deficient mice and control mice to CS for 8 weeks and compared cytokine production across lung, BALF and serum. CS exposure led to a disruption in cytokine production where cytokines like Il16 and Gcsf were dysregulated following CS exposure in the mutant mouse at a local and systemic level (**Fig.4**). Other cytokines like Mip2, Fractalkine and Il1a displayed differential expression in the *lincRNA-Cox2* mutant mouse following CS exposure suggesting that *lincRNA-Cox2* can impact some genes in a cell type specific manner. Mip2, Mip3b and Vegf were all found to be downregulated within the lungs of the *lincRNA-Cox2* deficient mice and interestingly they were all rescued back to WT levels in the *lincRNA-Cox2* MutxTg cross, indicating that *lincRNA-Cox2* can exert regulation on their production/secretion *in trans* (**Fig.4**).

Smoke exposure has been shown to lead to an exacerbated inflammatory response when a stimulus is introduced (52, 53); where CS-exposed mice displayed increased pulmonary inflammation and lung damage as well as increased production of some key inflammatory cytokines. CS exposure has also been shown to alter the ability of macrophages to respond to inflammatory stimuli (54). We observed that following 4.5 weeks exposure to CS the baseline levels of cytokines produced from BMDMs are similar between all genotypes and room air (RA) mice (**SFig. 8**). Interestingly, 8 weeks of CS exposure impacted the baseline levels of cytokines in BMDMs compared to RA suggesting that prolonged exposure to CS over time can impact the baseline expression of genes emerging from the bone marrow niche (**Fig. 1**). Genome wide RNA sequencing in combination with cytokine analysis revealed a skewed inflammatory response to LPS within smoke-exposed macrophages represented by dysregulation in coding genes expression and inflammatory cytokine secretion as well as an overall repression of lncRNA genes (**Fig.5**). This is especially interesting because it links CS exposure to a distorted immune response in which a macrophage has an altered response to a challenge, implicating CS in shaping the system’s response to infections. This is supported by mounting evidence that implicates CS exposure in subverting immunity where CS results in an attenuated immune response in which the host fails to respond effectively to a bacterial or viral challenge (55,56). Interestingly, loss of *lincRNA-Cox2* led to an exacerbated inflammatory response in LPS activated CS exposed macrophages, indicating that *lincRNA-Cox2* can shape the macrophage inflammatory response in CS exposed cells and alter their function.

In conclusion our work sheds light on the broad impact of smoking on immune responses in the lung, serum and the bone marrow niche. We uncovered a novel impact of CS on gene expression and splicing in bone marrow derived macrophages. In addition, we uncovered a novel role for *lincRNA-Cox2* in regulating immune responses to smoke. It is important for us to broaden our understanding of the mechanism that underlies inflammation if we are to identify and develop effective areas for therapeutic intervention for conditions such as COPD in the future.

## Materials and Methods

### Mice

Wild-type (WT) C57BL/6 mice were purchased from the Jackson Laboratory (Bar Harbor, ME) and bred at the University of California, Santa Cruz (UCSC). All mouse strains, including *lincRNA-Cox2* deficient (mutant or Mut) and MutxTg mice, were maintained under specific pathogen-free conditions in the animal facilities of UCSC and protocols performed in accordance with the guidelines set forth by UCSC and Weill Cornell Medical College Institutional Animal Care and Use Committees.

### Smoke Exposure Protocol

We selected age- and sex-matched mice, starting at 6–12 weeks of age, at random and exposed them to total body CS in a stainless-steel chamber using a whole-body smoke exposure device (Model TE-10 Teague Enterprises) for 2 hours per day, 5 days per week for 4.5 weeks or 8 weeks. Age-matched male and female mice were used for all CS exposures. Mice were exposed to CS in modular chambers as previously described (29,30). Briefly, using a TE-10 inhalation exposure apparatus (Teague Enterprises), mice were exposed to CS (100 3R4F cigarettes, University of Kentucky Center for Tobacco Reference Products) with an average total particulate matter (TPM) of 150 mg/m^3^ for 2 hours per day, 5 days per week for 4.5-8 weeks. For animals subjected to cigarette smoke exposure, early death was used as an exclusion criterion. At the end of the exposure regimen, we euthanized mice by CO2 narcosis, cannulated the tracheas and inflated the lungs with PBS at 25 cm of H2O pressure. eft lungs were tied off with a suture, dissected and placed into liquid nitrogen. Right lungs were extracted and fixed in 4% formalin at 4°C overnight.

### Cell culture

BMDMs were generated by culturing erythrocyte-depleted BM cells in DMEM supplemented with 10% FCS, 5 mL pen/strep (100×), 500 μL ciprofloxacin (10 mg/mL), and 10% L929 supernatant for 7 to 14 d, with the replacement of culture medium every 2 to 3 d.

### BMDM stimulations

Cells were stimulated with 200ng/mL LPS for the duration of 6hrs for RNA extraction and 18hrs for supernatant collection and secreted cytokine analysis.

### RNA isolation, cDNA synthesis and RT-qPCR

Total RNA was purified from cells or tissues using Direct-zol RNA MiniPrep Kit (Zymo Research, R2072) and TRIzol reagent (Ambion, T9424) according to the manufacturer’s instructions. RNA was quantified and assessed for purity using a nanodrop spectrometer (Thermo Fisher). Equal amounts of RNA (500 to 1,000 ng) were reverse transcribed using iScript Reverse Transcription Supermix (Bio-Rad, 1708841), followed by qPCR using iQ SYBR Green Supermix reagent (Bio-Rad, 1725122) with the following parameters: 50 °C for 2 min and 95 °C for 2 min, followed by 40 cycles of 95 °C for 15 s, 60 °C for 30 s, and 72 °C for 45 s, followed by melt-curve analysis to control for nonspecific PCR amplifications. Oligos used in qPCR analysis were designed using Primer3 Input version 0.4.0.

Gene expression levels were normalized to Gapdh or Hprt as housekeeping genes.

### PCR

RT-PCR validation was completed using three biological replicates. KAPA HiFi HotStart ReadyMix PCR Kit (Kapa Biosystems) and the manufacturer’s suggested cycling protocol were used to complete the PCR reaction with the following primers: Mse_Mib2_F1: AGGTGGACACCAAGAACCAG, Mse_Mib2_R1: GGATGCATGGTGTAGCAGTG. Yielding bands at length 548 bps and 452 bps for the inclusion and exclusion forms respectively. Band intensities were measured for each band in each condition and sample using ImageJ (57). The relative abundance of each isoform was calculated using the equation to calculate percent spliced in (PSI) PSI = inclusion counts/(inclusion counts + exclusion counts) in each condition and sample to validate the computationally derived delta PSI values.

### Cells Supernatant Collection for ELISA

Supernatant was collected from cultured and treated BMDMs, centrifuged at 12000xg, 5mins at RT and submitted for cytokine analysis.

### Serum Harvest

Mice were humanely sacrificed; blood was collected immediately postmortem by cardiac puncture. Blood was allowed to clot and centrifuged, serum was stored at −70°C, then sent to EVE for measurements of cytokines/chemokines.

### Lung Tissue Harvesting for cytokine measurement

Mice were humanely sacrificed, and their lungs were excised. The whole lungs were snap frozen and homogenized, and the resulting homogenates were incubated on ice for 30 min and then centrifuged at 300 × *g* for 20 min. The supernatants were harvested, passed through a 0.45-μm-pore-size filter, and used immediately or stored at −70°C, then sent to EVE for measurements of cytokines/chemokines.

### Harvesting Bronchoalveolar Lavage Fluid (BALF)

Bronchoalveolar Lavage Fluid (BALF) was harvested as previously stated by Cloonan et al. (29). Briefly, 40 mice were euthanized via CO2 asphyxiation, intubated with a 20G 3 1” catheter (SR-OX2025CA; Terumo), and the lungs were lavaged with ice cold PBS (10010-023; Life Technologies) supplemented with 0.5 mM EDTA (351-027-061; Quality Biological) in 0.5-ml increments for a total 4mLs. The BAL fluid (BALF) was collected from the first 0.5 ml after centrifugation at 500 3 g for 5 min at 4 °C and used for ELISA. The cell pellet was resuspended in 500 μl PBS, and leukocytes were counted using a hemocytometer. Specifically, 10 μl was removed for cell counting (performed in triplicate) using a hemocytometer. Cells were plated in sterile 12 well plates at 5e5/well (total of 8 wells) and use complete DMEM with 25 ng/ml supGM-CSF.

### RNA sequencing libraries

RNA-Seq was performed in monocytes, MDMs, and MDDCs. The data are accessible at the National Center for Biotechnology Information (NCBI) Gene Expression Omnibus (GEO) database, accession GSE184571.

RNA-Seq was performed in biological triplicates in WT RA and CS BMDMs at 0 and 6 h after LPS treatment (200 ng/mL). RNA-Seq libraries were generated from total RNA (1 μg) using the Bioo kit, quality was assessed, and samples were read on a High-SEq 4000 as paired-end 150-bp reads. Sequencing reads were aligned to the mouse genome (assembly GRCm38/mm10) using STAR. Differential gene-expression analyses were conducted using DESeq2. GO enrichment analysis was performed using PANTHER. Data was submitted to GEO, accession GSE184571.

### Alternative Splicing Analysis

Aligned reads (BAMs) were analyzed for alternative splicing events with rMATS v4.1.1 and JuncBASE (from Docker image mgmarin/juncbase:0.9). A custom GTF file combining long and short read sequencing data, and documented previously in Robinson *et al*. (13), was used by the programs as the input annotation. Following statistical analysis, the results from both analyses were filtered based on FDR, percent splice in (PSI), and number of reads supporting the event. Events that passed the following criteria are reported: FDR < 0.1, |PSI| > 10. Common intron retention events were identified using the intersect function of Bedtools v2.27.1. The Mib2 sashimi plot (Fig. 3C) was generated with rmats2sashimiplot v2.0.4. Parameters used for identifying common events using Bedtools intersect are: -wa -u -f -r 0.85 -a <rMATS intron retention events bed> -b <JuncBASE intron retention events bed>.

### rMATS

For rMATS analysis both the counting and statistical analysis were performed by the rMATS v4.1.1 package (58). In addition to filtering by FDR and PSI as described above, events were required to have total read support > 10, i.e. at least 10 reads supporting the inclusion or exclusion event across all samples. The following parameters were used -t paired --readLength 100 --gtf <custom.gtf> --libType fr-unstranded --allow-clipping.

### JuncBASE + DRIMSeq

In this analysis JuncBASE was used for generating event counts and DRIMSeq v1.20 was used for determining statistically significant events (59,60). An intron coordinate file (BED format) was downloaded from the UCSC Table Browser with Gencode vM18 as for use in JuncBASE. The JuncBASE event counts were reformatted for DRIMSeq analysis with an inhouse script. Following statistical analysis, the results from both analyses were filtered based on FDR and percent splice in (PSI). Parameters used for JuncBASE were -j <intron coordinates from Gencode vM18> --jcn_seq_len 188. Parameters used for finding significantly differentially spliced events using DRIMSeq were: min_samps_gene_expr = 6, min_samps_feature_expr = 3, min_gene_expr = 10, min_feature_expr = 0.

## Supporting information

SupFig.1

SupFig.2

SupFig.3

SupFig.4

SupFig.5

SupFig.6

SupFig.7

SupFig.8

## Author Contributions

SC and SMC conceptualized and acquired funding for this study. EP, AC, KK and WZZ performed cigarette-smoke animal model and harvested all samples for study. EKR and MMS generated all BMDMs and performed RNA-sequencing for this study. EM performed RNA-sequencing and alternative splicing analysis for this study. MMS and EKR both wrote the initial draft of this manuscript. All authors contributed to reviewing and editing of manuscript.

## Acknowledgements

This work was supported by the California Tobacco Related Disease Research Program grant 27IP-0017 to S.C and the Cornelius Hopper diversity award from the TRDRP to M.M.S.

## Figure Legends

**Supplementary Figure 1: Cigarette smoke COPD does not globally regulate the transcriptional expression of inflammatory transcription factors**. (A) Bar plot of Log2FcMAP determined by DESeq2 analysis of BMDMs from room air *vs*. cigarette smoked mice. Asterisks indicate statistically significant differences between mouse lines using DESeq2 (*p < 0.05).

**Supplementary Figure 2: Gene ontology analysis of CS down-regulated protein-coding genes in BMDMs**. (A) The associated biological process of down-regulated differentially expressed protein-coding genes from CS exposure using DAVID tools.

**Supplementary Figure 3: Computational pipeline and comparison of t-test and DRIMSeq alternative splicing events**.

(A) Bioinformatic pipeline for human and mouse RNA-seq data. Alternative splicing event-type classification of significant differential splicing events (|ΔPSI| > 10 and adjusted p-value < 0.1) in mouse macrophages ± cigarette smoke (CS) exposure.

**Supplementary Figure 4: Cigarette Smoke induces alternative splicing in long non-coding RNA bone marrow-derived macrophages**. (A) Volcano plot of all differentially expressed long non-coding RNA when comparing BMDMs from room air (RA) to cigarette smoke (CS) mice with significantly differentially spliced genes (FDR < 0.1, |ΔPSI| > 10) across at least one event type marked.

**Supplementary Figure 5: Mib2 is not differentially expressed in bone marrow-derived macrophages from cigarette smoke exposed mice**. (A) The normalized counts and DESeq2 analysis of *Mib2*.

**Supplementary Figure 6: Full PCR gel image of Mib2 retained intron event across samples**(A) PCR gel results of Mib2 at the RI event site and loading control HPRT in biological triplicates. Lanes 1 and 14 are 100 bp ladder. Lanes 2 through 4 are RA samples 1,2,3 and lanes 5 through 7 are CS samples 1,2,3 showing PCR amplification of the Mib2 RI event. When the intron is retained a band of size 548 bps is expected, otherwise a band of size 452 bps. Lanes 8 through 13 are RA samples 1,2,3 followed by CS samples 1,2,3 showing amplification of the housekeeping gene Hprt.

**Supplementary Figure 7: Changes in inflammatory cytokine levels between WT and lincRNA-Cox2 mutant mice under RA and CS treatments for 8 weeks**. ELISAs were performed on serum for (B) Epo and (C) Fractalkine. ELISAs were performed on normalized lung tissue homogenates on (D) Mip3b, (E) Vegf, (F) Ip10, (G) Kc, (H) Mcp1 and (I) Timp1. Each dot represents an individual animal. Error bars represent standard deviation of biological replicates. Asterisks indicate statistically significant differences between mouse lines using Student’s t-tests (*p < 0.05, **p < 0.01, ***p < 0.001). Student’s t tests were performed using GraphPad Prism to obtain p values.

**Supplementary Figure 8: lincRNA-Cox2 does not regulate cytokines in BMDMs from cigarette smoked mice**.

(A) Experimental schematic for BMDM harvested from WT, lincRNA-Cox2 mutant and lincRNA-Cox2 MutxTg mice exposed to room air or cigarette-smoked. Supernatants were harvested and ELISAs were performed to measure (B) Gcsf, (C) Il16, (D) IL1a, (E) Il11, (F) IP10, (G) Fractalkine, (H) MCSF, (I) MDC and (J) MIP-2. Each dot represents an individual animal. Error bars represent standard deviation of biological triplicates. Asterisks indicate statistically significant differences between mouse lines using Student’s t-tests (*p < 0.05, **p < 0.01). Student’s t tests were performed using GraphPad Prism to obtain p values.

