## Supplementary figures and images for "*LincRNA-Cox2* regulates smoke-induced inflammation in murine macrophages"

### SupFig.1

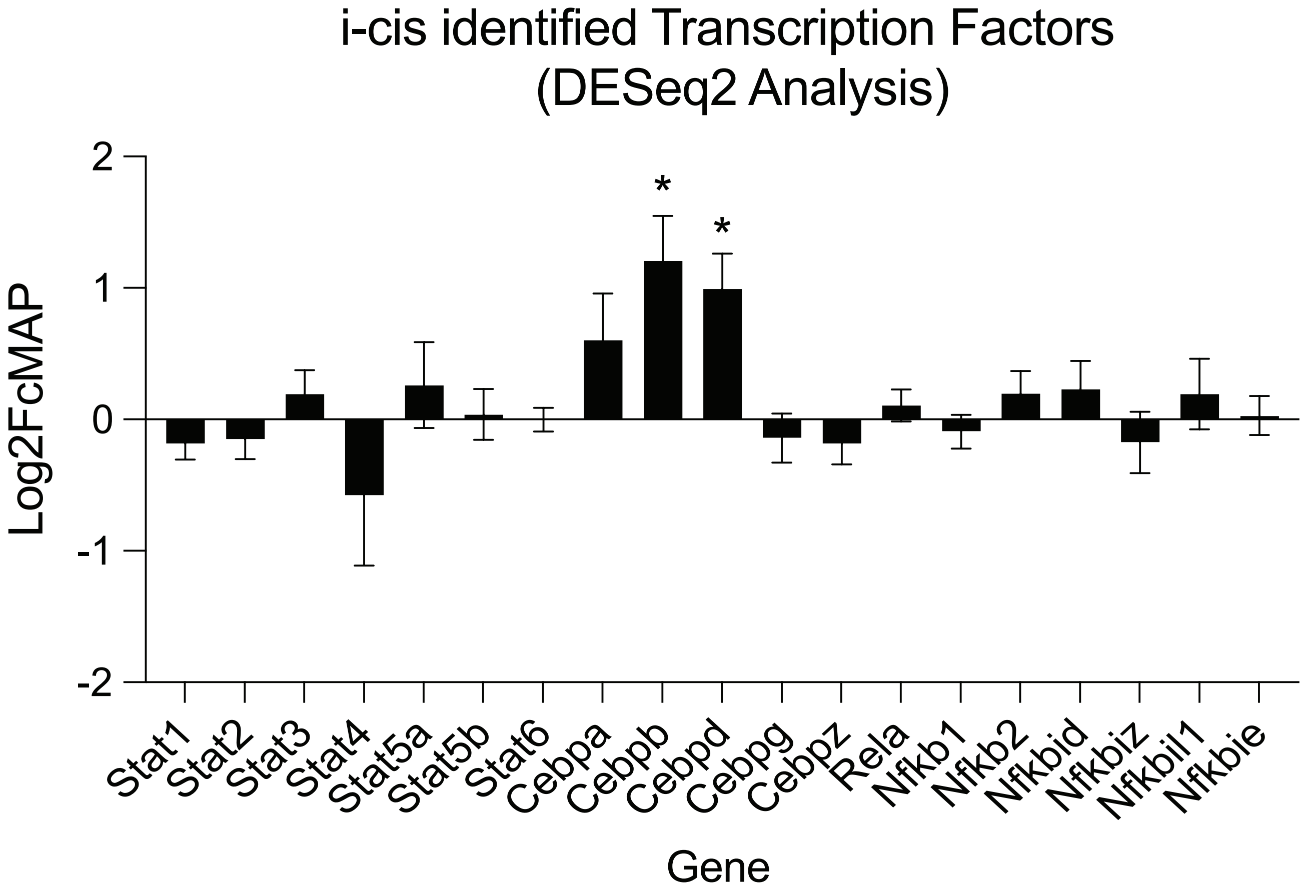

### SupFig.2

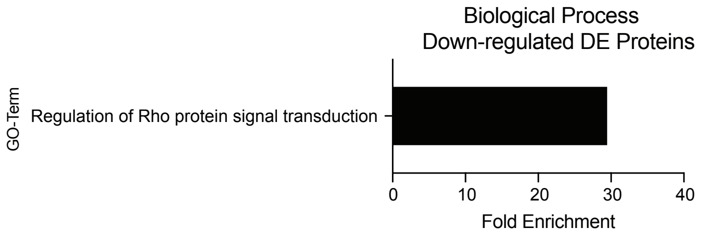

### SupFig.3

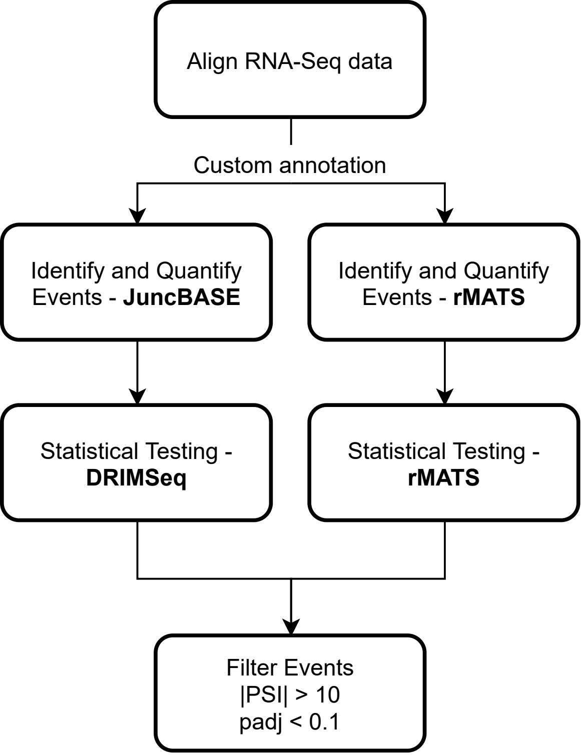

### SupFig.4

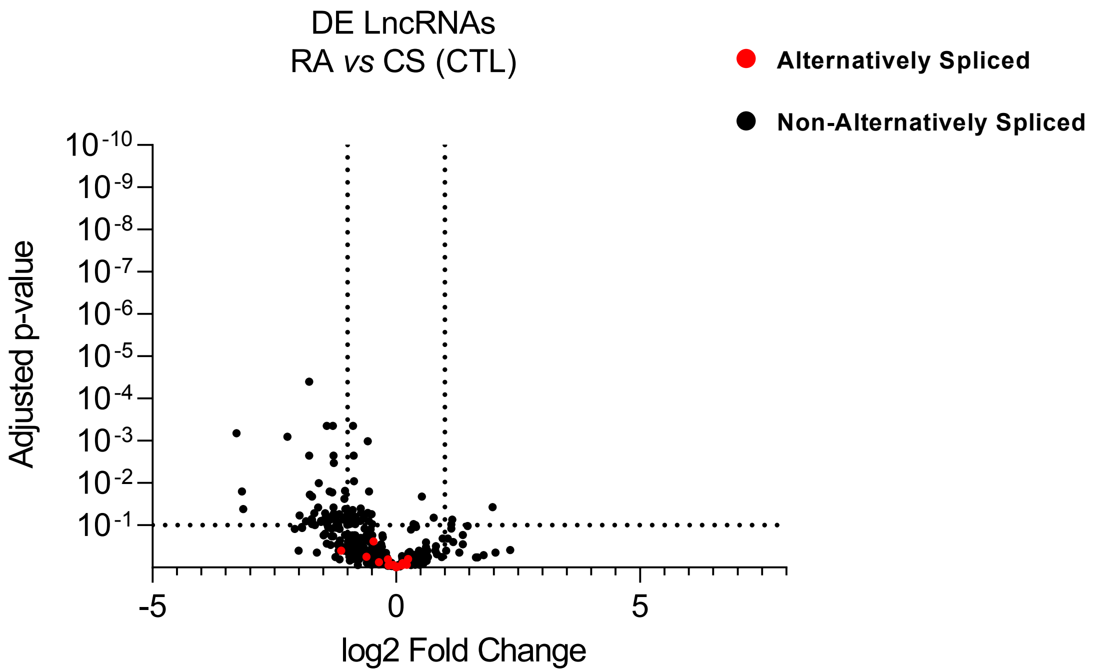

### SupFig.5

A.

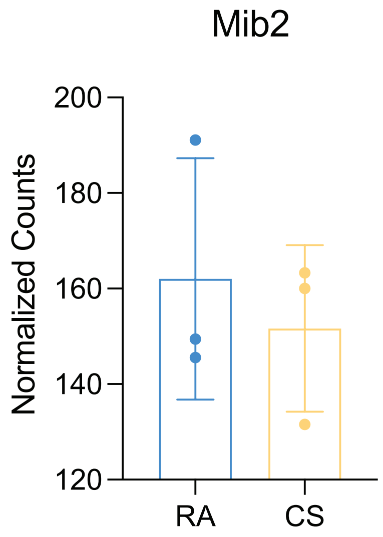

### SupFig.6

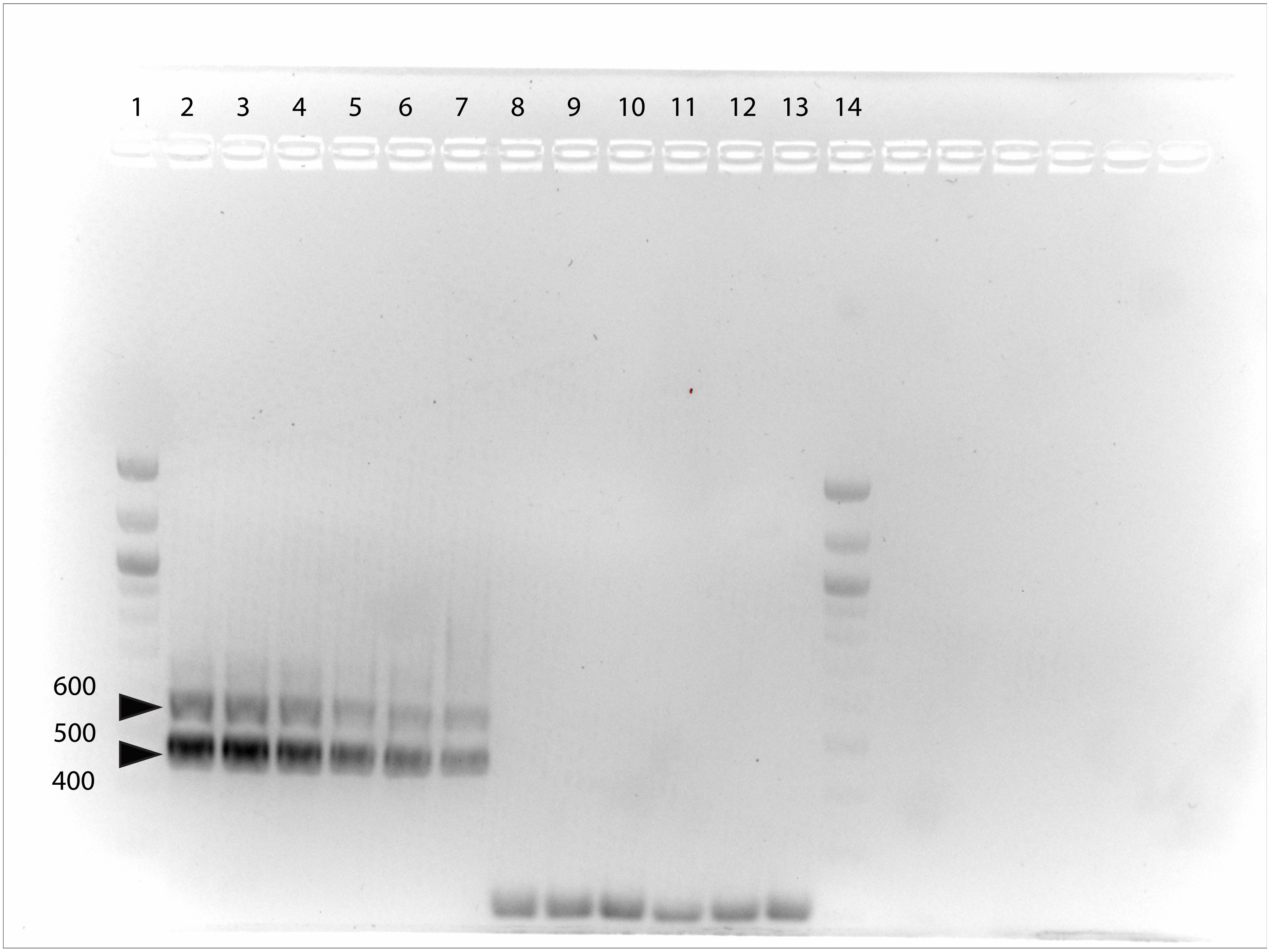

### SupFig.7

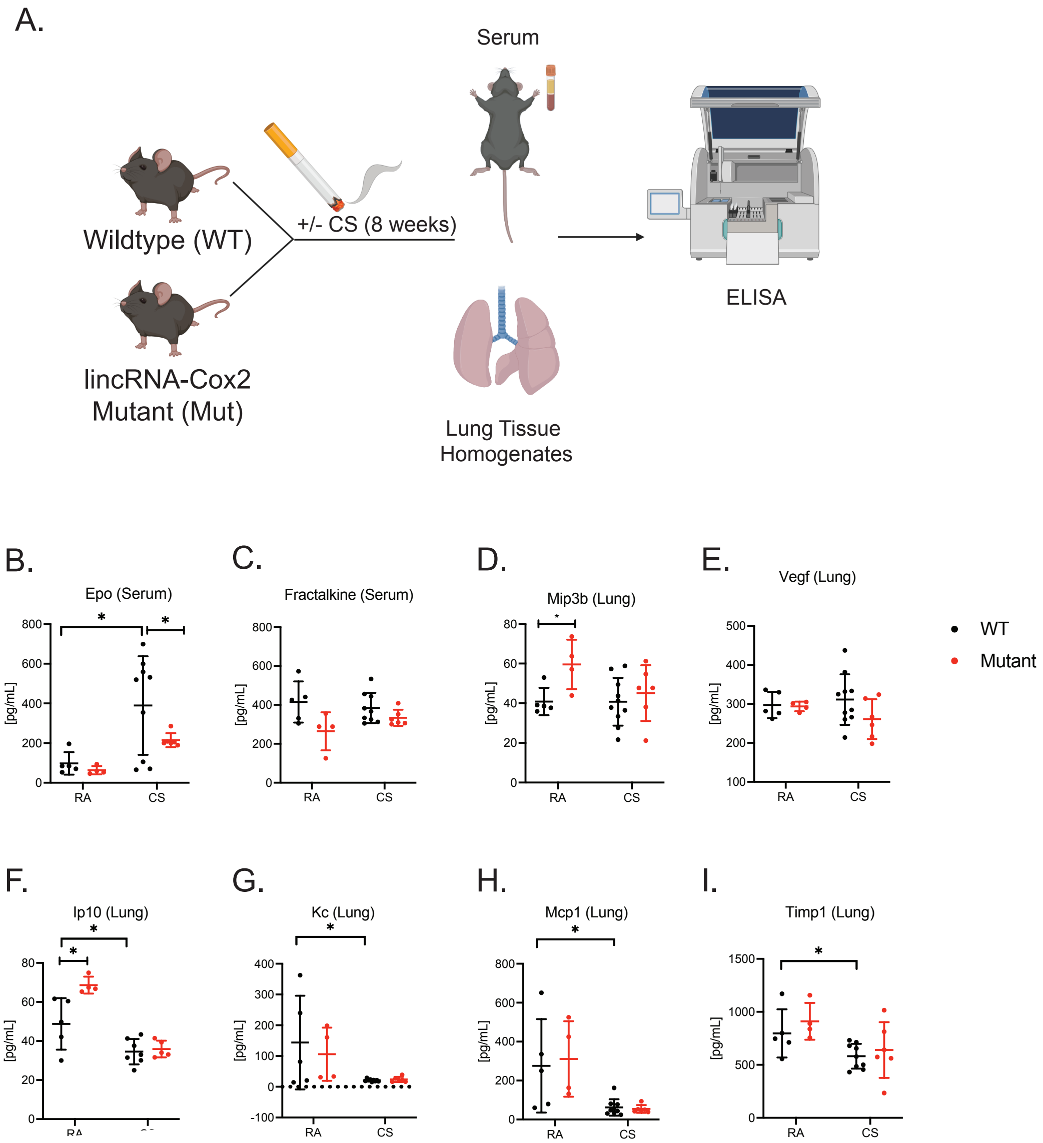

### SupFig.8

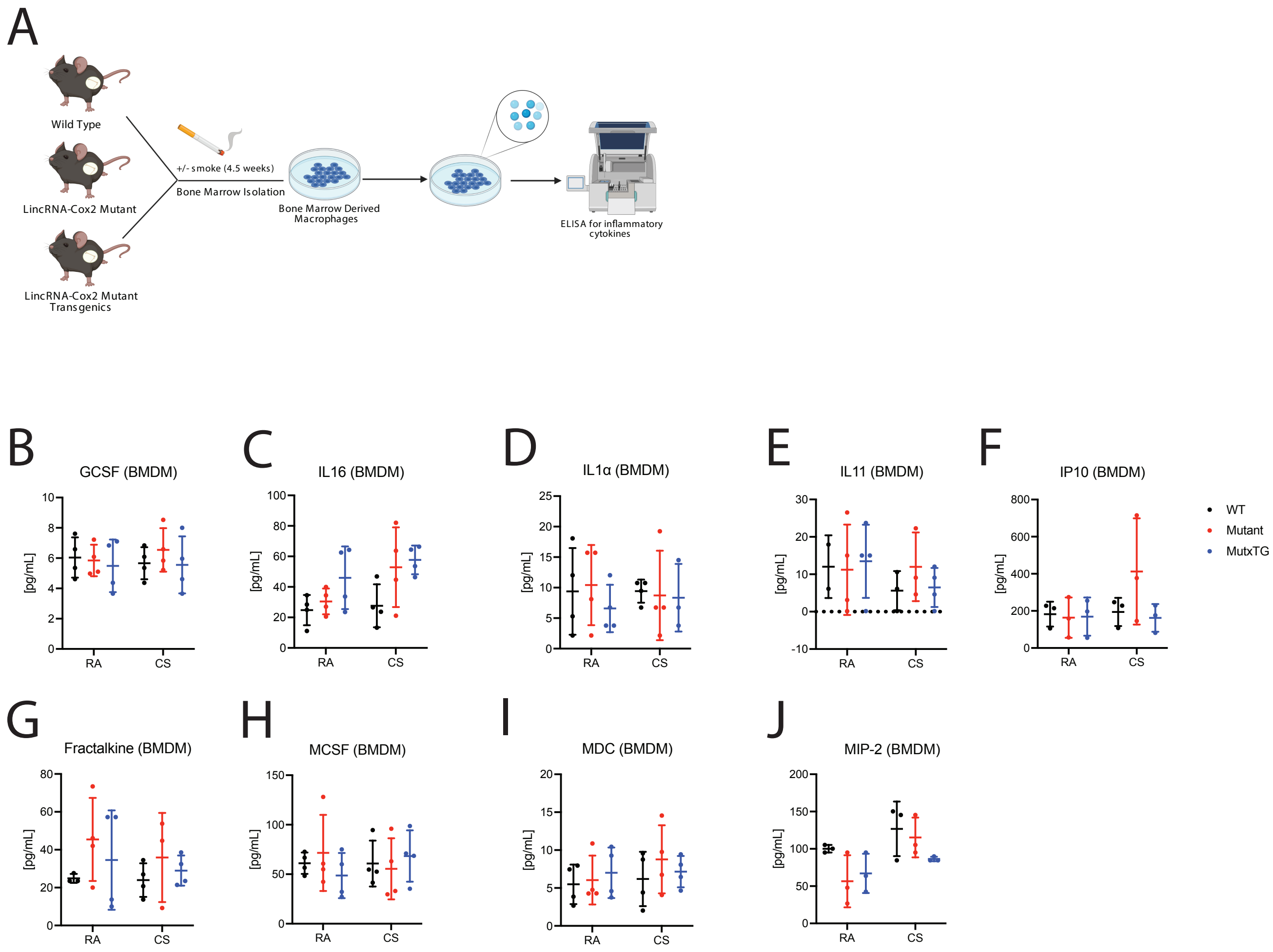
